# The *Arabidopsis* ERF transcription factor ORA59 coordinates jasmonic acid- and ethylene-responsive gene expression to regulate plant immunity

**DOI:** 10.1101/2021.03.16.435681

**Authors:** Young Nam Yang, Youngsung Kim, Hyeri Kim, Su Jin Kim, Kwang-Moon Cho, Yerin Kim, Dong Sook Lee, Myoung-Hoon Lee, Soo Young Kim, Jong Chan Hong, Sun Jae Kwon, Jungmin Choi, Ohkmae K. Park

**Affiliations:** Department of Life Sciences, Korea University, Seoul 02841, Korea; Molecular Diagnosis Division, AccuGene, Incheon 22006, Korea; Department of Biomedical Sciences, Korea University College of Medicine, Seoul 02841, Korea; Department of Biotechnology and Kumho Life Science Laboratory, College of Agriculture and Life Sciences, Chonnam National University, Gwangju 61186, Korea; Division of Life Science, Plant Molecular Biology and Biotechnology Research Center, Gyeongsang National University, Jinju 52828, Korea; Animal and Plant Quarantine Agency, Foot and Mouth Disease Research Division, Gimcheon 39660, Korea; Genuine Research, Seoul 06040, Korea

**Keywords:** *Arabidopsis*, ORA59, transcription factor, ERELEE4, GCC box, Ethylene, jasmonic acid, plant immunity

## Abstract

Jasmonic acid (JA) and ethylene (ET) signaling modulate plant defense against necrotrophic pathogens. These hormone pathways lead to transcriptional reprogramming, which is a major part of plant immunity and requires the roles of transcription factors. ET response factors are responsible for the transcriptional regulation of JA/ET-responsive defense genes, among which ORA59 functions as a key regulator of this process and has been implicated in the JA-ET crosstalk. Here, we identified the ERELEE4 as an ORA59-binding *cis*-element, in addition to the well-characterized GCC box, demonstrating that ORA59 regulates JA/ET-responsive genes through direct binding to these elements in the gene promoters. Notably, ORA59 exhibited differential preference for the GCC box and ERELEE4, depending on whether ORA59 activation is achieved by JA and ET, respectively. Our results provide insights into how ORA59 can generate specific patterns of gene expression dynamics through JA and ET hormone pathways.

## Introduction

In nature, plants encounter a wide range of microbial pathogens with varying lifestyles and infection strategies. Upon pathogen recognition, plants rapidly activate defense responses, and the levels of resistance are influenced by hormone actions (De Vos et al., 2005; Pieterse et al., 2009). Salicylic acid (SA), jasmonic acid (JA), and ethylene (ET) are primary defense hormones that trigger immune signaling mechanisms (Dong, 1998; Pieterse et al., 2012). Classically, SA signaling enhances resistance against biotrophic and hemibiotrophic pathogens such as *Hyaloperonospora arabidopsidis* and *Pseudomonas syringae*, whereas JA and ET signaling activate resistance against necrotrophic pathogens such as *Alternaria brassicicola* and *Botrytis cinerea* (Feys and Parker, 2000; Glazebrook, 2005). Antagonism between SA and JA/ET and synergism between JA and ET have been mostly observed in studies of plant immunity, although there is evidence of positive interactions between them (Kim et al., 2013; Koornneef and Pieterse, 2008; Penninckx et al., 1998; Thomma et al., 1998). These hormone signaling pathways are interconnected in a complex network and their crosstalk enables plants to tailor defense responses efficiently (Bostock, 2005; Kunkel and Brooks, 2002; Spoel and Dong, 2008).

JA and ET modulate diverse developmental processes and defense responses in plants (Broekgaarden et al., 2015; Joo and Kim, 2007; Zhu and Lee, 2015). Their signaling pathways work by de-repression mechanisms. MYC2/3/4 transcription factors play essential roles in JA signaling, and in the absence of JA, remain in repressed states by binding to transcriptional repressors jasmonate ZIM-domain (JAZ) proteins (Chini et al., 2007; Thines et al., 2007). JA promotes the interaction between JAZs and the F-box protein coronatine insensitive 1 (COI1), resulting in degradation of JAZs and de-repression of MYCs (Katsir et al., 2008; Sheard et al., 2010; Yan et al., 2009). The activated MYCs then regulate gene expression and various JA responses (Cheng et al., 2011; Fernández-Calvo et al., 2011; Niu et al., 2011). ET insensitive 2 (EIN2) and EIN3 are key positive regulators of ET signaling (Alonso et al., 1999; Chao et al., 1997). In the absence of ET, ET receptors activate the Raf-like serine/threonine kinase constitutive triple response 1 (CTR1), which phosphorylates EIN2 to repress its activity (Kieber et al., 1993). EIN3 and its homolog EIN3-like 1 (EIL1) are also targeted for degradation by EIN3-binding F-box protein 1 (EBF1) and EBF2 (Guo and Ecker, 2003; Potuschak et al., 2003). ET binding to ET receptors deactivates CTR1, which is followed by de-repression of EIN2 (Chao et al., 1997). In the activation process, EIN2 is cleaved and its C-terminal fragment translocates into the nucleus and inhibits EBF1/2, promoting EIN3/EIL1 accumulation (Ju et al., 2012; Qiao et al., 2012; Wen et al., 2012). EIN3/EIL1 further activate downstream genes, including the ET response factor (ERF) family transcription factors (Chang et al., 2013; Solano et al., 1998). EIN3/EIL1 and ERFs regulate ET-mediated gene expression. Plant defense against necrotrophic pathogens requires JA and ET, and synergistic and interdependent interactions between JA and ET have been described (Koornneef and Pieterse, 2008; Thomma et al., 1998). ERFs are important regulators of the JA-ET crosstalk, and in particular, ERF1 and octadecanoid-responsive arabidopsis 59 (ORA59), belonging to the group IX ERF family, have been suggested to act as integrators of ET and JA signaling (Lorenzo et al., 2003; Pré et al., 2008b). The expression of *pathogenesis-related* (*PR*) genes such as *plant defensin 1.2* (*PDF1.2*) and *basic chitinase* (*b-CHI*) was synergistically induced by JA and ET, and abolished in JA-insensitive *coi1* and ET-insensitive *ein2* mutants, while depending on ERF1 and ORA59 (Lorenzo et al., 2003; Penninckx et al., 1998; Pré et al., 2008b). Analysis of the *PDF1.2* promoter indicated that ERF1 and ORA59 induce *PDF1.2* expression through direct binding to GCC boxes in the *PDF1.2* promoter (Zarei et al., 2011). Like other *PR* genes, the expression of *ERF1* and *OAR59* themselves exhibited a synergistic response to JA and ET, which was impaired in *coi1* and *ein2* mutants. The role of ERF1 and ORA59 in defense has been revealed in *ERF1*- and *ORA59*-overexpressing plants displaying enhanced resistance to necrotrophic pathogens (Berrocal-Lobo et al., 2002; Pré et al., 2008b; Kim et al., 2018).

*ERF1* and *ORA59* have been determined to be regulated by EIN3 and their JA- and ET-responsive expression was abolished in *ein3 eil1* mutant (Solano et al., 1998; Zander et al., 2012; Zhu et al., 2011). Given that EIN3 is a positive regulator of *ERF1* and *ORA59*, it was assessed whether EIN3 and EIL1 control JA and ET synergy on defense gene expression. JAZ proteins recruited histone deacetylase 6 (HDA6) as a corepressor to deacetylate histones and interacted with EIN3/EIL1 to repress EIN3/EIL1-mediated transcription (Solano et al., 1998; Zander et al., 2012; Zhu et al., 2011). JA led to JAZ degradation and removed JAZ-HDA6 from EIN3/EIL1, and ET enhanced EIN3/EIL1 accumulation, enabling EIN3/EIL1 to converge JA and ET signaling. The role as an integrative hub for JA and ET signaling has also been assigned to subunits of the Mediator complex that connects transcription factors with the RNA polymerase II machinery (Bäckström et al., 2007). The Mediator subunit MED25 physically interacted with several transcription factors, including ERF1, ORA59, and EIN3/EIL1, and was required for ERF1- and ORA59-activated *PDF1.2* expression (Çevik et al., 2012; Yang et al., 2014). On the other hand, SA suppressed JA-dependent transcription by negatively affecting ORA59 protein abundance, suggesting that ORA59 acts as a node for SA and JA antagonism (He et al., 2017; Van der Does et al., 2013).

In this study, we report that the previously undefined *cis*-element ERELEE4 is critical for JA/ET-induced transcription and is frequently present in the promoters of JA/ET-responsive genes. In a yeast one-hybrid (Y1H) screening, ORA59 was identified as a specific transcription factor that binds to the ERELEE4 element, although ORA59 was previously known to regulate gene transcription by binding to the GCC box. Depending on whether plants are exposed to JA or the ET precursor 1-aminocyclopropane-1-carboxylic acid (ACC) and ET, ORA59 exhibited preferential binding to the GCC box and ERELEE4, respectively. The present study explores how two defense hormones JA and ET coordinate gene expression required for plant immunity through the regulation of ORA59.

## Results

### ERE is the *cis*-acting element for ET-responsive *GLIP1* expression

We previously demonstrated that *Arabidopsis GDSL lipase 1* (*GLIP1*) is an ET-responsive defense gene that confers resistance to necrotrophic pathogens (Kim et al., 2014; Kim et al., 2013; Kwon et al., 2009; Oh et al., 2005). In an expression analysis, Col-0 plants exhibited strong *GLIP1* expression in response to ACC and ET, but a slight increase in *GLIP1* expression upon methyl JA (MeJA) treatment (Supplementary Fig. 1a,b), which is in line with previous results (Kim et al., 2013; Oh et al., 2005). However, both ACC- and MeJA-induced *GLIP1* expression was abolished in ET-insensitive *ein2* and *ein3 eil1* and JA-insensitive *coi1* mutants, indicating that *GLIP1* induction requires both ET and JA signaling pathways (Supplementary Fig. 1c,d). Whereas EIN2- and EIN3/EIL1-dependent *GLIP1* expression is consistent with our previous observation (Kim et al., 2013), COI1 dependency may result from EIN3 regulation by COI1-JAZ (Zhu et al., 2011).

To investigate how *GLIP1* expression is modulated by ET, the *GLIP1* promoter was searched for the *cis*-element critical for ET-responsive *GLIP1* expression. In previous studies, promoter analysis of ET- and JA-responsive *PR* genes led to the identification of the conserved sequence AGCCGCC or GCC box that serves as a binding site for ERFs (Brown et al., 2003; Hao et al., 1998; Ohme-Takagi and Shinshi, 1995). Accordingly, we expected the presence of the GCC box in the *GLIP1* promoter and scanned the 2966-bp *GLIP1* promoter region upstream of the transcription start site for *cis*-acting elements, using the PLACE program (http://www.dna.affrc.go.jp/PLACE/). This analysis revealed that the *GLIP1* promoter has no GCC box sequences and is enriched with binding motifs related to hormone and pathogen responses, which include two ET-responsive elements, ERELEE4 (AWTTCAAA) and RAV1AAT (CAACA) (Supplementary Table 1). ERELEE4 has been identified in promoter regions of tomato *E4* and carnation *glutathione-S-transferase 1* (*GST1*) genes, but poorly characterized (Itzhaki et al., 1994; Montgomery et al., 1993). RAV1AAT has been isolated as the binding motif for the *Arabidopsis* related to ABI3/VP1 1 (RAV1) transcription factor belonging to the APETALA2/ERF superfamily (Kagaya et al., 1999). ERELEE4 was located at 4 positions and RAV1AAT at 11 positions, here designated as ERE1 to ERE4 and RAV1 to RAV11, respectively, upward from the transcription start site. In the case of EREs, there were two different sequences, A<u>T</u>TTCAAA at ERE1, ERE3, and ERE4 and A<u>A</u>TTCAAA at ERE2.

To examine whether ERE and RAV are key regulatory elements for ET-induced *GLIP1* expression, we introduced the chimeric constructs of the *GLIP1* promoter (*pGLIP1*) and the *β-glucuronidase* (*GUS*) reporter gene into *Arabidopsis* protoplasts and performed transient GUS reporter assays. In accordance with previous results (Kim et al., 2013), *pGLIP1* elevated GUS activity in response to ET and ACC, compared to mock treatments (Fig. 1a and Supplementary Fig. 2a). *pGLIP1* constructs with a series of 5′ deletions (*pGLIP1*−2466, −1466, −966, and −566) were made and assayed for their ability to drive ET/ACC-induced GUS expression. As *pGLIP1* became shorter and RAV and ERE elements were lost, GUS activity decreased proportionally. No GUS activity was driven by *pGLIP1*-966 containing 3 RAV elements (RAV1 to RAV3). The ability of ERE and RAV to respond to ET/ACC was further tested using synthetic promoters, in which a minimal promoter (TATA-box) was fused to four tandem copies (4x) of ERE and RAV and their mutated versions mERE and mRAV (Fig. 1b and Supplementary Fig. 2b). Among two ERE sequences in *pGLIP1*, more frequent ATTTCAAA was used for the synthetic promoter. The 4xERE promoter strongly triggered ET/ACC-induced GUS expression compared to 4xRAV, suggesting that ERE is critical for ET/ACC-mediated *GLIP1* expression. Their mutated versions had little effect on GUS expression. Next, *pGLIP1*-mediated GUS activity was measured after mutation of one by one or 4 EREs at once (Fig. 1c). *pGLIP1* with individual ERE mutations displayed significantly decreased GUS activity (47-69% reduction) compared to the native promoter. GUS activity driven by *pGLIP1^mEREs^* with all 4 EREs mutated was largely eliminated. *pGLIP1* activity was further assessed in Col-0 plants harboring *pGLIP1:GUS* or *pGLIP1^mEREs^:GUS* reporters. Histochemical staining developed strong GUS signals in *pGLIP1:GUS* plants upon ACC treatment and in response to *B. cinerea* infection, which were largely abolished in *pGLIP1^mEREs^:GUS* plants (Fig. 1d). These results together demonstrate that ERE plays a major role in ET-responsive *GLIP1* expression.

**Figure 1.**
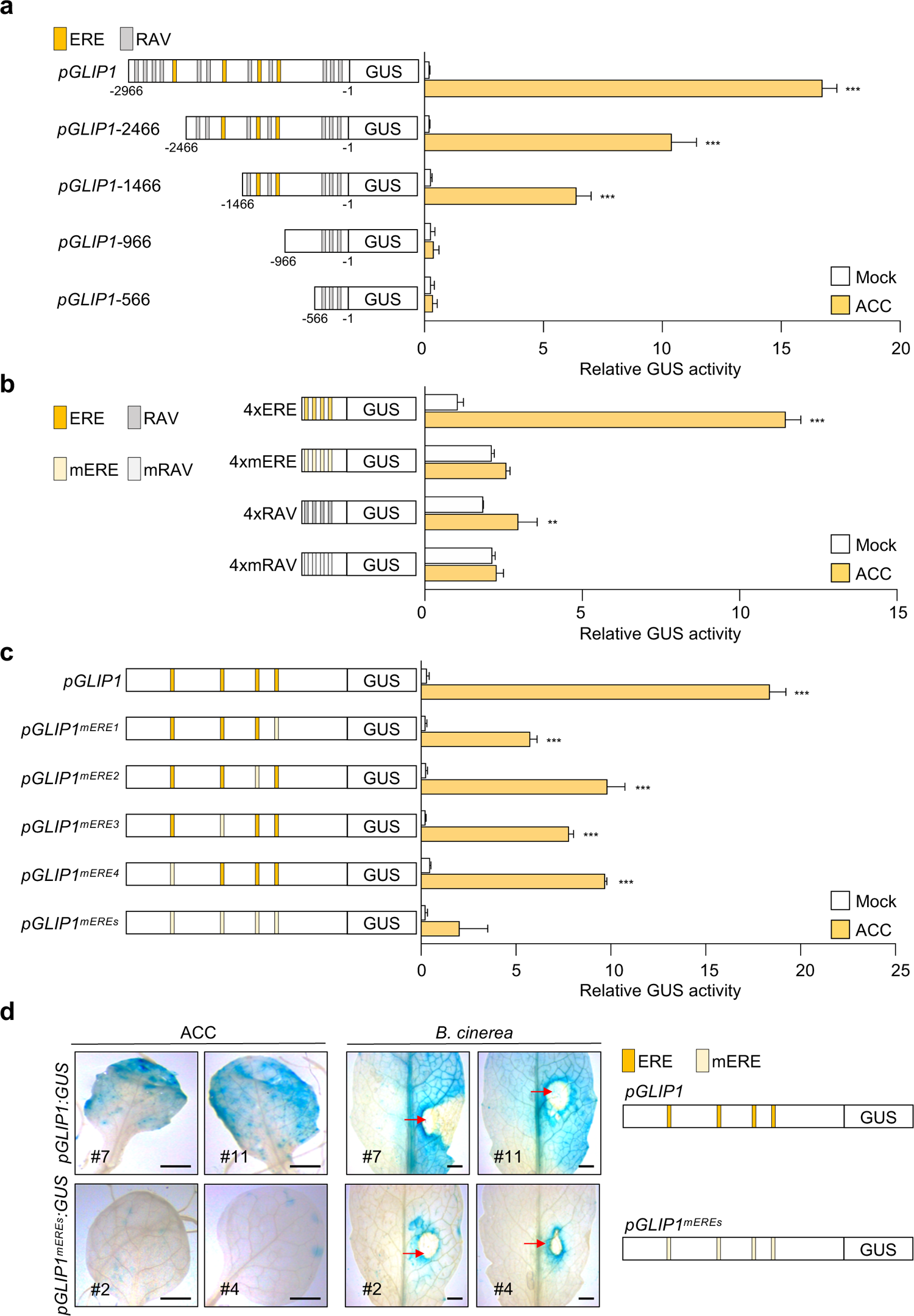
ERE is the essential regulatory element in the *GLIP1* promoter. **a**, GUS reporter assays showing ACC-induced expression of the *GUS* reporter gene driven by the full-length and truncated *GLIP1* promoters. The left panel illustrates deletions of the *GLIP1* promoter. The ERE and RAV elements in the *GLIP1* promoter are boxed in yellow and gray, respectively. **b**, GUS reporter assays showing ACC-induced expression of the *GUS* reporter gene driven by synthetic promoters of 4xERE and 4xRAV, and their mutant versions 4xmERE and 4xmRAV. The left panel illustrates synthetic promoters. **c**, GUS reporter assays showing ACC-induced expression of the *GUS* reporter gene driven by the *GLIP1* promoters with individual or all ERE mutations. The left panel illustrates ERE mutations of the *GLIP1* promoter. **d**, GUS staining of ACC- and *B. cinerea*-treated leaves of transgenic plants expressing the *GUS* reporter gene driven by native or ERE-mutated *GLIP1* promoters. Six-week-old *pGLIP1:GUS* and *pGLIP1^mEREs^:GUS* plants were treated with ACC (1 mM) for 6 h or with 5 µl droplets of *B. cinerea* spore suspensions (5 x 10^5^ spores ml^-1^) for 2 days. Representative images are provided, and infection sites are indicated by red arrows. Bars, 1 mm. In **a**-**c**, transfected protoplasts were treated with mock (water) and ACC (200 μM) for 6 h. Values represent means ± SD (*n* = 3 biological replicates). Asterisks indicate significant differences from mock treatment as determined by one-way ANOVA with Tukey test (\*\**P* < 0.01; \*\*\**P* < 0.001).

The requirement of ERE elements for *GLIP1* expression was additionally assessed by generating transgenic plants, *pGLIP1:GLIP1-GFP* and *pGLIP1^mEREs^:GLIP1-GFP*, which express GLIP1 fused to green fluorescent protein (GFP) at the C-terminus under the control of *pGLIP1* and *pGLIP1^mEREs^*, respectively, in the *glip1* mutant background. First, plants were infected with *A. brasscicola*, and examined for disease development. Whereas *glip1* mutant was highly susceptible to *A. brasscicola*, *pGLIP1*-driven *GLIP1-GFP* expression restored disease resistance in *glip1* (Fig. 2a-c). Consistently, *GLIP1-GFP* transcripts and GLIP1-GFP proteins accumulated and GFP fluorescence was detected in *pGLIP1:GLIP1-GFP* plants, but not in *pGLIP1^mEREs^:GLIP1-GFP* plants, in response to *A. brasscicola* and ACC treatments (Fig. 2d-f). These results together indicate that ERE elements are essential for *GLIP1* expression during the immune response.

**Figure 2.**
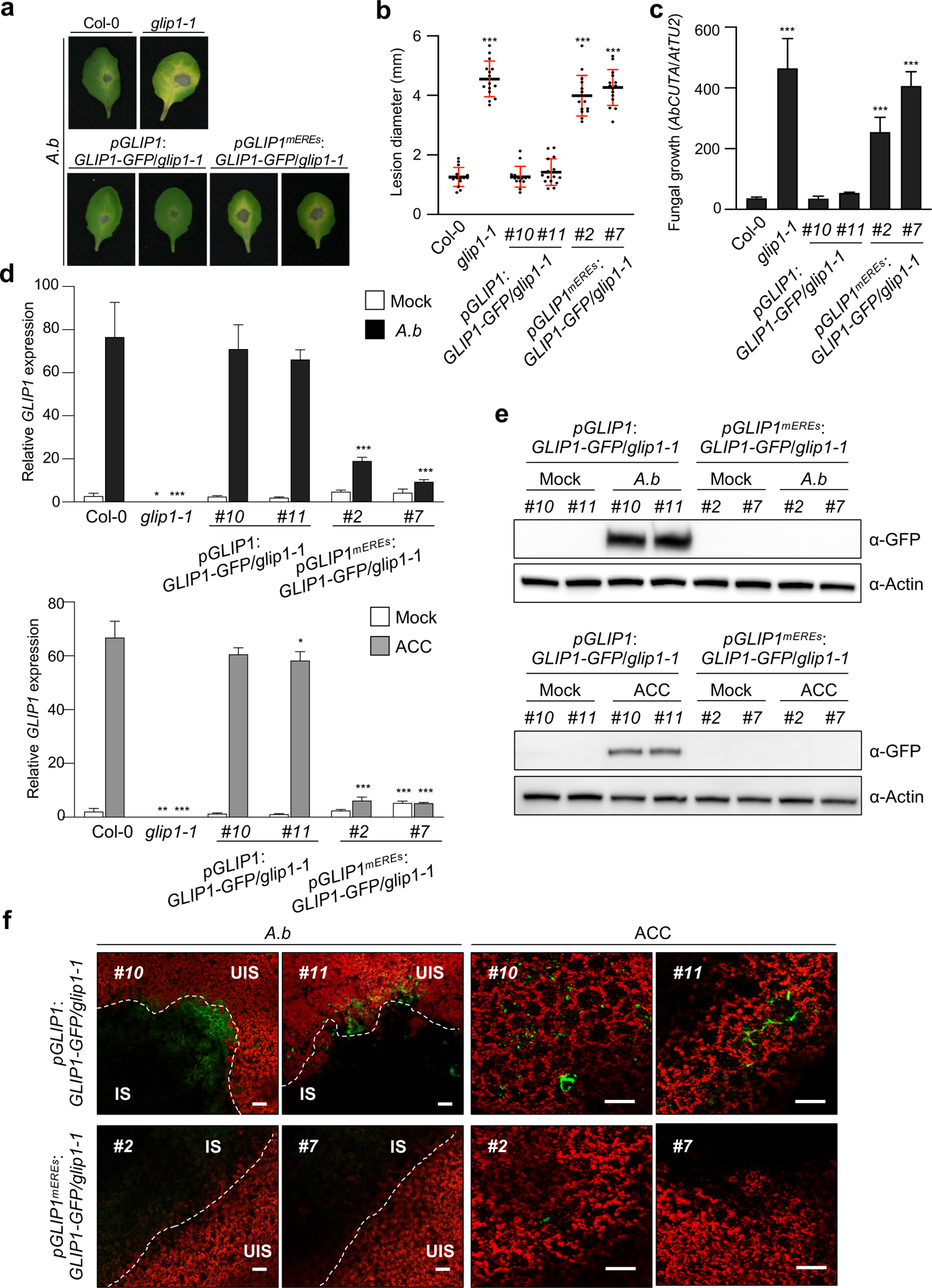
ERE is required for *GLIP1* expression during the immune response. **a**,**b**, Disease symptoms (**a**) and lesion diameters (**b**) in leaves inoculated with *A. brassicicola*. Values represent means ± SD (*n* = 15 infected leaves). **c**, Measurement of *A. brassicicola* growth in infected leaves. The abundance of *A.brassicicola cutinase A* (*AbCUTA*) gene relative to *Arabidopsis tubulin 2* (*AtTU2*) was analyzed by qPCR. Values represent means ± SD (*n* = 6 infected leaves). **d**, Analysis of *GLIP1* expression in *A. brassicicola*- and ACC-treated plants. Values represent means ± SD (*n* = 3 biological replicates). **e**, Immunoblot analysis of GLIP1-GFP expression in *A. brassicicola*- and ACC-treated plants. Protein extracts were subjected to immunoblotting with anti-GFP and anti-Actin antibodies. Actin levels served as a control. **f**, Confocal images of GLIP1-GFP expression in *A. brassicicola*- and ACC-treated plants. Bars, 100 μm. Two independent transgenic lines were used in all experiments. Six-week-old plants were treated with ACC (1 mM) for 6 h (**d**-**f**) or with 5 µl droplets of *B. cinerea* spore suspensions (5 x 10^5^ spores ml^-1^) for 1 (**d**) and 2 (**a**,**b**,**c**,**e**,**f**) days. In **b**-**d**, asterisks indicate significant differences from the respective Col-0 as determined by one-way ANOVA with Tukey test (\**P* < 0.05; \*\**P* < 0.01; \*\*\**P* < 0.001). IS, infected site; UIS, uninfected site; *A.b*, *A. brassicicola*.

### ORA59 is an ERE-binding transcription factor

Next, we searched for a transcription factor(s) that regulate ET-responsive *GLIP1* expression, and for this, performed Y1H screening using the ERE sequence ATTTCAAA as bait. The yeast strain, harboring three tandem copies of ERE fused to *HIS3* and *lacZ* genes, was transformed with a prey library composed of 1050 *Arabidopsis* transcription factor cDNAs (Welchen et al., 2009). Screening of 2 x 10^6^ transformants yielded 84 positive clones growing on selective media lacking histidine and containing 3-amino-1,2,4-triazole (3-AT) (Supplementary Table 2). Among these positive clones, ORA59, related to AP2.2 (RAP2.2), and caprice-like MYB3 (CPL3) were subjected to further analysis, as they most strongly increased β-galactosidase reporter activity. Re-transformation with the recovered plasmid DNAs enabled yeast cells to grow on selective media (Fig. 3a).

**Figure 3.**
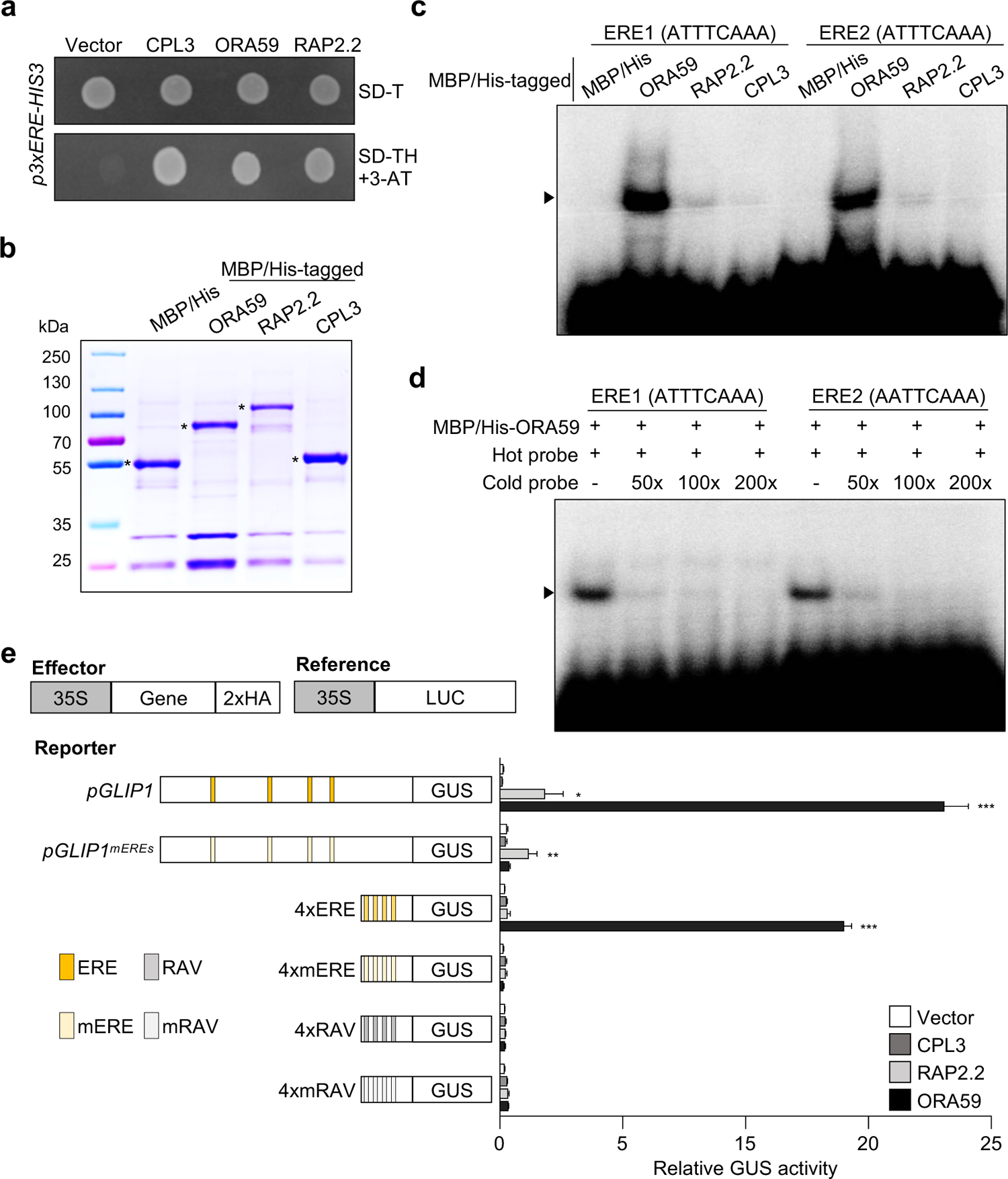
ORA59 is the specific ERE-binding transcription factor. **a**, Isolation of ERE-binding transcription factors by Y1H screening. Yeast cells harboring the 3xERE-*HIS3* reporter gene were transformed with effector constructs *CPL3*, *ORA59*, and *RAP2.2*. Transcription factor binding to the ERE was tested on a selective medium lacking tryptophan and histidine, and supplemented with 0.2 mM 3-amino-1,2,4-triazole (SD-TH + 3-AT). **b**, Coomassie blue staining of purified recombinant ORA59, RAP2.2, and CPL3 fused with N-terminal MBP and His tags. Asterisks indicate the corresponding purified proteins. **c**, DNA binding of ORA59, RAP2.2, and CPL3 to the two ERE sequences ATTTCAAA (ERE1) and AATTCAAA (ERE2). Recombinant proteins were incubated with biotin-labeled ERE oligonucleotide probes in EMSA. **d**, Competition assays. ORA59 binding to the ERE1/2 was competed with increasing amounts (50x, 100x, 200x) of unlabeled oligonucleotide competitors. **e**, Transactivation analysis showing the ORA59-mediated *GUS* reporter gene induction driven by the *GLIP1* and 4xERE synthetic promoters. The left panel illustrates reference, effector, and reporter constructs. Reporter DNAs, either alone or together with effector DNAs, were transfected into protoplasts, and GUS activity was measured. Luciferase (LUC) expressed under control of the CaMV 35S promoter was used as an internal control (reference). Values represent means ± SD (*n* = 3 biological replicates). Asterisks indicate significant differences from vector control as determined by one-way ANOVA with Tukey test (\**P* < 0.05; \*\**P* < 0.01; \*\*\**P* < 0.001). MBP, maltose-binding protein; 35S, CaMV 35S; 2xHA, two copies of the hemagglutinin (HA) tag sequence.

To test for *in vitro* binding of ORA59, RAP2.2, and CPL3 to the ERE element, we performed an electrophoretic mobility shift assay (EMSA) using recombinant proteins and DNA probes of two ERE sequences ATTTCAAA and AATTCAAA (Fig. 3b,c). Whereas ORA59 formed a shifted band, RAP2.2 and CPL3 exhibited weak binding. ORA59 had similar levels of binding activity to these two ERE sequences. The addition of excess amounts of unlabeled ERE probes effectively competed the binding, verifying specific ORA59 binding to the ERE sequences (Fig. 3d). Transient GUS reporter assays were then performed to determine whether they can induce transcription through the ERE *in vivo* (Fig. 3e). The *pGLIP1* and synthetic 4xERE and 4xRAV promoters, and their mutant versions *pGLIP1^mEREs^*, 4xmERE, and 4xmRAV were used as reporter constructs, and together with effector constructs of CPL3, RAP2.2, and ORA59, were transformed into *Arabidopsis* protoplasts. Whereas ORA59 strongly activated *pGLIP1*- and 4xERE-mediated GUS expression, slight GUS expression driven by both *pGLIP1* and *pGLIP1^mEREs^* was observed with RAP2.2, and no GUS activity was observed with CPL3. Transactivation by ORA59 was dependent on the ERE in reporter genes, because no activity was detected for reporters with *pGLIP1^mEREs^* and 4xmERE or with 4xRAV and 4xmRAV. These results suggest that ORA59 controls *GLIP1* expression via ERE binding. Because the GCC box has been determined as a specific binding site for ORA59 and other ERFs in previous reports (Hao et al., 1998; Ohme-Takagi and Shinshi, 1995; Zarei et al., 2011), the binding activity of ORA59 to GCC box and ERE was compared using recombinant ORA59 proteins. In addition to full-length ORA59, several truncated forms of ORA59 were prepared (Fig. 4a-c). In the EMSA analysis, full-length ORA59 bound to GCC box more strongly than to ERE (Fig. 4d). Noticeably, N-terminal deletion (F1) dramatically enhanced ORA59 binding to GCC box, but rather abrogated ERE-binding activity. In contrast, ORA59 with partial C-terminal deletion (F3) showed much stronger ERE binding. We failed to secure soluble ORA59 proteins with deletion of the entire C-terminal region after the AP2 domain. The C-terminal (F2) and N-terminal (F4) portions alone exhibited no DNA binding activities, as expected for ORA59 without the DNA-binding AP2 domain. AP2 domain alone had stronger binding activity to ERE than to GCC box, which was reversed for full-length ORA59. These results suggest that ORA59 may form distinct structural conformations in binding to ERE and GCC box, and the N-terminal and C-terminal portions of ORA59 affect ORA59 binding to ERE and GCC box in different manners. The N-terminal portion may have a positive or negative effect on ORA59 binding to ERE and GCC box, respectively, and this may be the opposite for the C-terminal portion.

**Figure 4.**
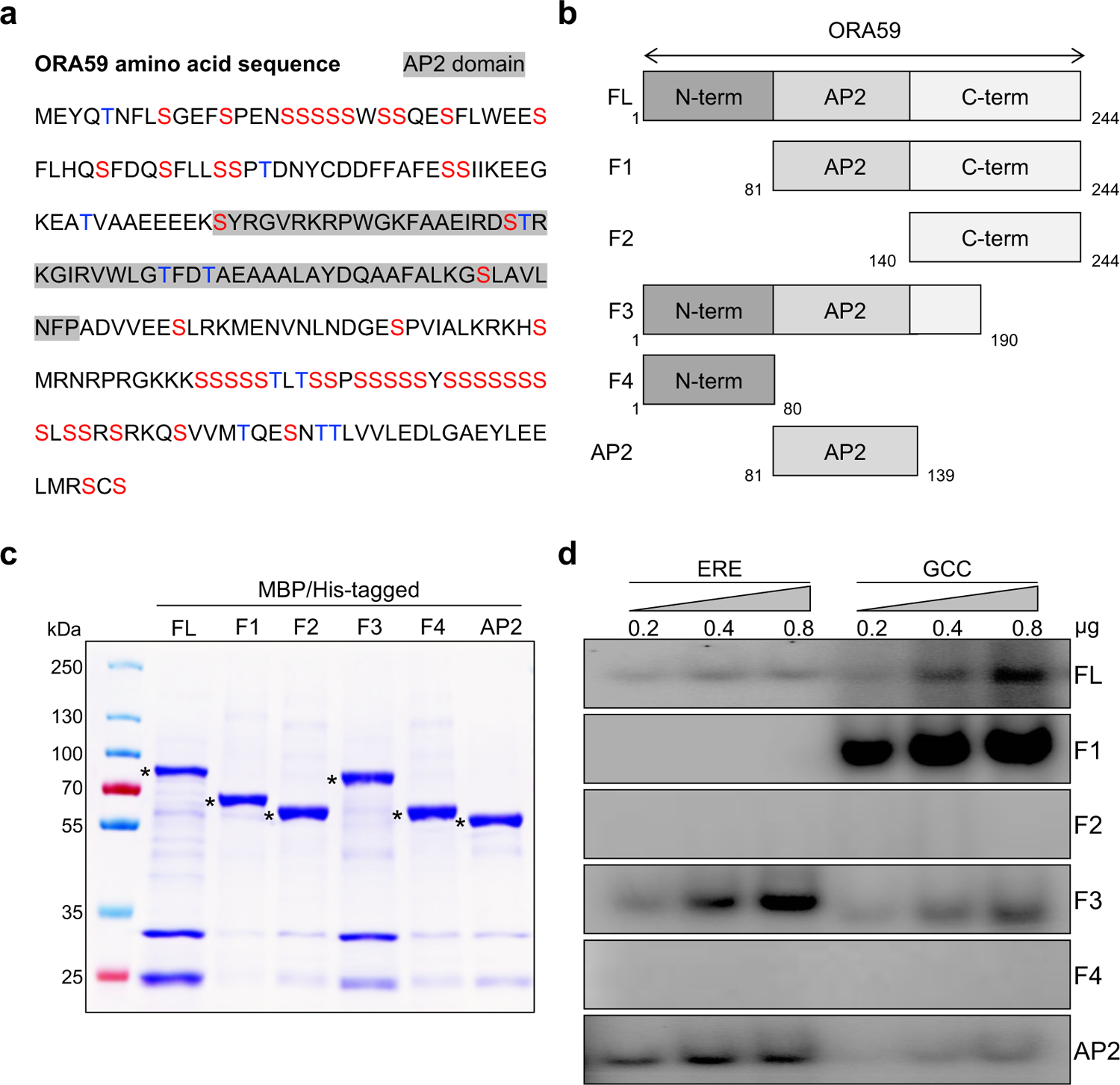
Truncated forms of ORA59 show distinct binding to ERE and GCC box. **a**, Amino acid sequence of ORA59 protein. Ser and Thr residues are indicated in red and blue, respectively. The AP2 domain is shaded in gray. **b**, Schematic diagram of full-length and truncated ORA59 proteins prepared for EMSA. **c**, Coomassie blue staining of purified recombinant full-length and truncated ORA59 fused with N-terminal MBP and His tags. Asterisks indicate the corresponding purified proteins. **d**, DNA binding of different forms of ORA59 to the ERE and GCC box elements. Increasing amounts (0.2, 0.4, and 0.8 μg) of recombinant proteins were incubated with biotin-labeled ERE and GCC box oligonucleotide probes in EMSA. FL, full-length; GCC, GCC box.

### ORA59 binding to ERE and GCC box is differentially regulated in ET and JA signaling

To study how ORA59 interacts with ERE and GCC box elements *in vivo*, we additionally prepared transgenic plants (*35S:ORA59-GFP*) overexpressing ORA59 fused to GFP at the C-terminus under the control of the CaMV 35S promoter (Supplementary Fig. 3a,b). As observed in a previous study (Pré et al., 2008b), *35S:ORA59-GFP* plants showed a dwarf phenotype. Basal transcript levels of *GLIP1* were increased in *35S:ORA59-GFP* plants, and ACC- and *B. cinerea*-induced *GLIP1* expression was diminished in *ora59* mutant, confirming that ORA59 is the key regulator of *GLIP1* expression (Supplementary Fig. 3c). *35S:ORA59-GFP* plants exhibited enhanced resistance against *B. cinerea*, as determined by lesion size and abundance of fungal actin gene (Supplementary Fig. 3d). In contrast, susceptibility to *B. cinerea* was increased in *ora59* plants. Among independent lines, *35S:ORA59-GFP* (*#6*) was used for further study.

ORA59-GFP protein levels were monitored in *35S:ORA59-GFP* plants exposed to ACC and MeJA. ORA59-GFP proteins rapidly disappeared in the presence of the protein synthesis inhibitor cycloheximide (CHX), which was repressed by treatment with the proteasome inhibitor MG132 (Fig. 5a). ORA59-GFP protein abundance was elevated in ACC- or MeJA-treated *35S:ORA59-GFP* plants. These results indicate that ORA59 undergoes 26S proteasome-dependent degradation, and ET and JA enhance the stability of ORA59 proteins.

**Figure 5.**
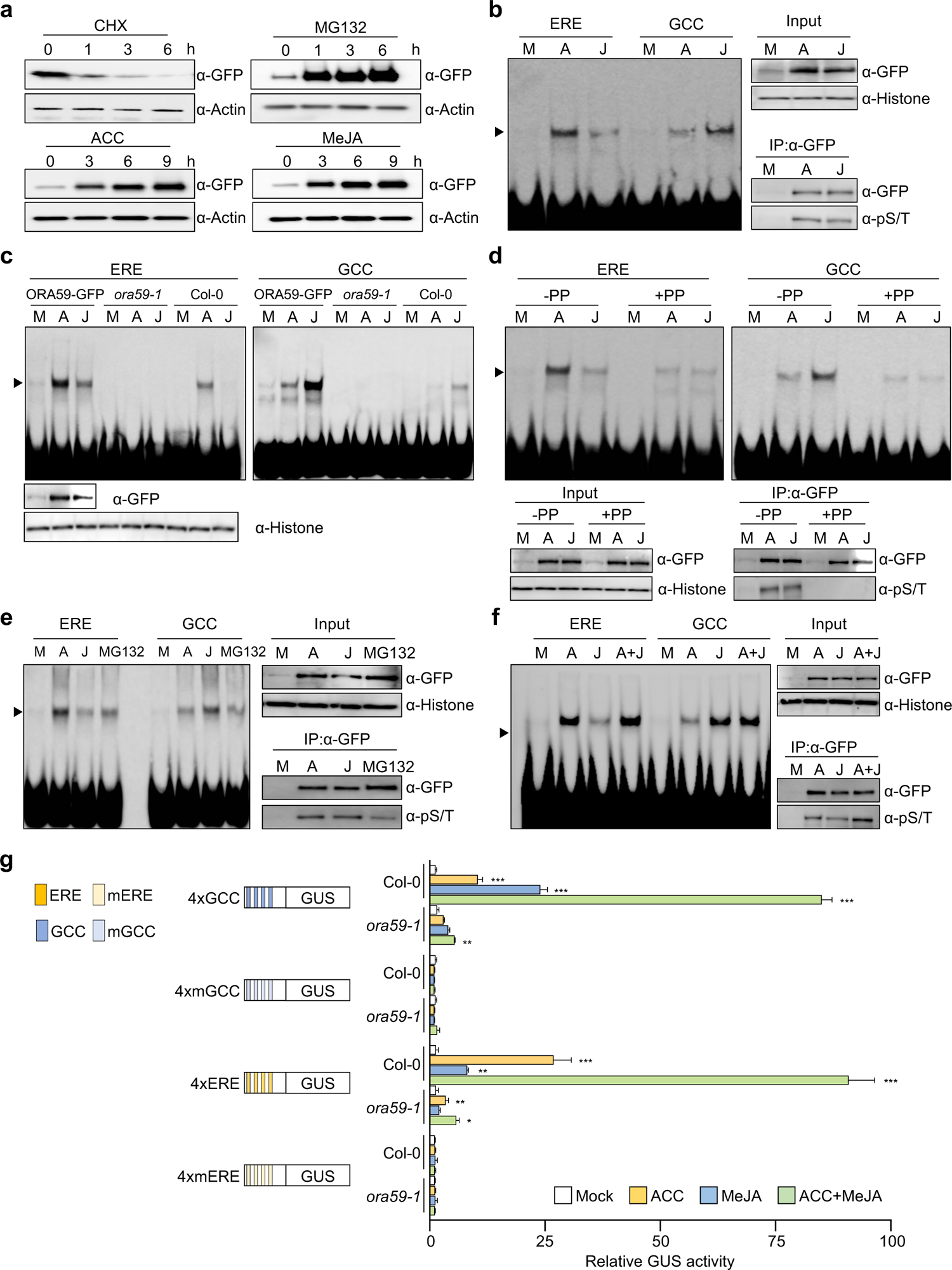
ORA59 binding to ERE and GCC box is regulated in ACC- and JA-dependent manners. **a**, Immunoblot analysis of ORA59 stability in *35S*:*ORA59*-*GFP* plants. Six-week-old plants were treated with cycloheximide (100 µM), MG132 (50 µM), ACC (1 mM), and MeJA (100 µM) for the indicated times. Protein extracts were subjected to immunoblotting with anti-GFP and anti-Actin antibodies. Actin levels served as a control. CHX, cycloheximide. **b**, EMSA analysis of nuclear extracts from *35S*:*ORA59*-*GFP* plants. Six-week-old plants were treated with ACC (1 mM) and MeJA (100 µM) for 6 h. Nuclear extracts were incubated with biotin-labeled ERE and GCC box oligonucleotide probes in EMSA. **c**, EMSA analysis of nuclear extracts from *35S*:*ORA59*-*GFP*, *ora59*, and Col-0 plants. Six-week-old plants were treated with ACC (1 mM) and MeJA (100 µM) for 6 h. Nuclear extracts were incubated with biotin-labeled ERE and GCC box oligonucleotide probes in EMSA. **d**, Effect of phosphatase treatment on DNA binding of ORA59. Six-week-old plants were treated with ACC (1 mM) and MeJA (100 µM) for 6 h. Nuclear extracts were treated with lambda phosphatase for 30 min before incubation with biotin-labeled ERE and GCC box oligonucleotide probes in EMSA. **e**, Effect of MG132 treatment on DNA binding of ORA59. Six-week-old plants were treated with ACC (1 mM), MeJA (100 µM), and MG132 (50 µM) for 6 h. Nuclear extracts were incubated with biotin-labeled ERE and GCC box oligonucleotide probes in EMSA. **f**, Effect of ACC and MeJA co-treatments on DNA binding of ORA59. Six-week-old plants were treated with ACC (1 mM), MeJA (100 µM), and a combination of ACC (1 mM) and MeJA (100 µM) for 6 h. Nuclear extracts were incubated with biotin-labeled ERE and GCC box oligonucleotide probes in EMSA. **g**, GUS reporter assays showing the effect of ACC and MeJA co-treatments on the expression of the *GUS* reporter gene driven by synthetic promoters of 4xERE and 4xRAV, and their mutant versions 4xmERE and 4xmRAV. The left panel illustrates synthetic promoters. Transfected Col-0 and *ora59* protoplasts were treated with mock (water), ACC (200 μM), MeJA (20 µM), and a combination of ACC (200 μM) and MeJA (20 µM) for 6 h. Values represent means ± SD (*n* = 3 biological replicates). Asterisks indicate significant differences from mock treatment as determined by one-way ANOVA with Tukey test (\**P* < 0.05; \*\**P* < 0.01; \*\*\**P* < 0.001). In **b**-**f**, ORA59 levels (input) in nuclear extracts were determined by immunoblotting with anti-GFP and anti-Histone H3 antibodies. Histone levels served as a control. In **b**,**d**-**f**, to assess the phosphorylation status of ORA59, nuclear extracts were incubated with an anti-GFP antibody and the immunoprecipitated ORA59-GFP proteins were subjected to immunoblotting with anti-GFP and anti-phospho-Ser/Thr (pS/T) antibodies. IP, immunoprecipitation; GCC, GCC box; PP, phosphatase; A, ACC; J, MeJA.

To examine ORA59 binding to ERE and GCC box *in planta*, nuclear extracts were prepared from ACC/ET- and MeJA-treated *35S:ORA59-GFP* plants and assessed for binding to these elements by EMSA. Noticeably, nuclear extracts from ACC/ET- and MeJA-treated *35S:ORA59-GFP* plants showed DNA binding activities with differential preference for ERE and GCC box, respectively, which were largely diminished in Col-0 and *ora59* extracts (Fig. 5b,c and Supplementary Fig. 4). These results imply that DNA-protein complexes observed are mostly of ORA59 expressed in *35S:ORA59-GFP* plants. Col-0 nuclear extracts, albeit weakly binding, retained the hormone-dependent preference for ERE and GCC box.

The Ser-rich sequence of ORA59 (Fig. 4a) led us to speculate that DNA binding properties of ORA59 may be regulated by post-translational modifications, such as phosphorylation. To address this, nuclear extracts of *35S:ORA59-GFP* plants were immunoprecipitated with an anti-GFP antibody, and the isolated proteins were subjected to Western blotting with an anti-phospho-Ser/Thr antibody. It revealed that ORA59 is phosphorylated in ACC- and MeJA-treated plants (Fig. 5b). To further verify this, *35S:ORA59-GFP* nuclear extracts were treated with lambda phosphatase before incubation with DNA probes. Phosphatase treatment led to dephosphorylation of ORA59, which was accompanied by a significant reduction in ORA59 binding to ERE and GCC box, and particularly, lack of hormone-dependent binding sequence specificity (Fig. 5d). When accumulated after MG132 treatment, ORA59-GFP proteins showed similar results with much lower level of phosphorylation, compared to those treated with ACC and MeJA (Fig. 5e). This indicates that ORA59 is normally phosphorylated to a certain extent and the phosphorylation level is increased in response to ET and JA. These results suggest that ET- and JA-mediated ORA59 phosphorylation is critical for ORA59 activity.

Considering the role of ORA59 in the ET-JA crosstalk, the next question was how ORA59 binds to ERE and GCC box when activated by two hormones simultaneously. To investigate this, *35S:ORA59-GFP* plants were co-treated with ACC and JA, and then subjected to EMSA. A combination of ACC and JA did not further increase DNA binding of ORA59, nor did it change ORA59 protein abundance, compared to treatment with each hormone (Fig. 5f). In contrast, the level of ORA59 phosphorylation was largely increased by ACC and MeJA co-treatments. We then examined whether hormone-dependent DNA binding properties of ORA59 are correlated with transcriptional activity. GUS reporter assays were conducted using 4xERE and 4xGCC box synthetic promoters. In Col-0 protoplasts, ACC treatment induced a large increase in transcription through the ERE but a less increase through the GCC box (Fig. 5g). Conversely, a large increase was observed with the GCC box but a modest increase with the ERE in response to MeJA treatment. Simultaneous treatments with ACC and MeJA led to a synergistic increase in GUS activity through both ERE and GCC box. This transcriptional activation was not observed with mutated elements and significantly decreased in *ora59* protoplasts. These results suggest that ET- and JA-regulated transcription is associated with differential DNA binding of ORA59, and ORA59 regulates ET and JA synergy at the level of transcriptional activation, but not at the level of DNA binding.

### ORA59 regulates gene expression by direct binding to ERE and GCC box

It was then determined whether other genes are also regulated by ORA59 in ET/JA-dependent ways. We found that *PDF1.2* genes have different distributions of ERE and GCC box in their promoters such that promoters of *PDF1.2a*, *PDF1.2b*, and *PDF1.2c* have a single GCC box, one GCC box and two ERE, and a single ERE elements, respectively (Fig. 6a). We conducted GUS reporter assays using *PDF1.2* promoters. Whereas GUS expression driven by *PDF1.2* promoters was induced by both ACC and MeJA, *PDF1.2a* with only GCC box and *PDF1.2c* with only ERE responded more strongly to MeJA and ACC, respectively, than to the other (Fig. 6b). Likewise, mutations of respective elements in *PDF1.2* promoters largely affected transcriptional activation, and in particular, mutation of either ERE or GCC box in the *PDF1.2b* promoter containing both elements more significantly reduced ACC- or MeJA-induced transcription, respectively. Gene expression analysis in Col-0 plants showed that endogenous transcript levels of *PDF1.2* genes were increased by ACC and MeJA treatments with similar preference for hormones observed in GUS reporter assays, and this increase was abolished in *ora59* plants (Fig. 6c).

**Figure 6.**
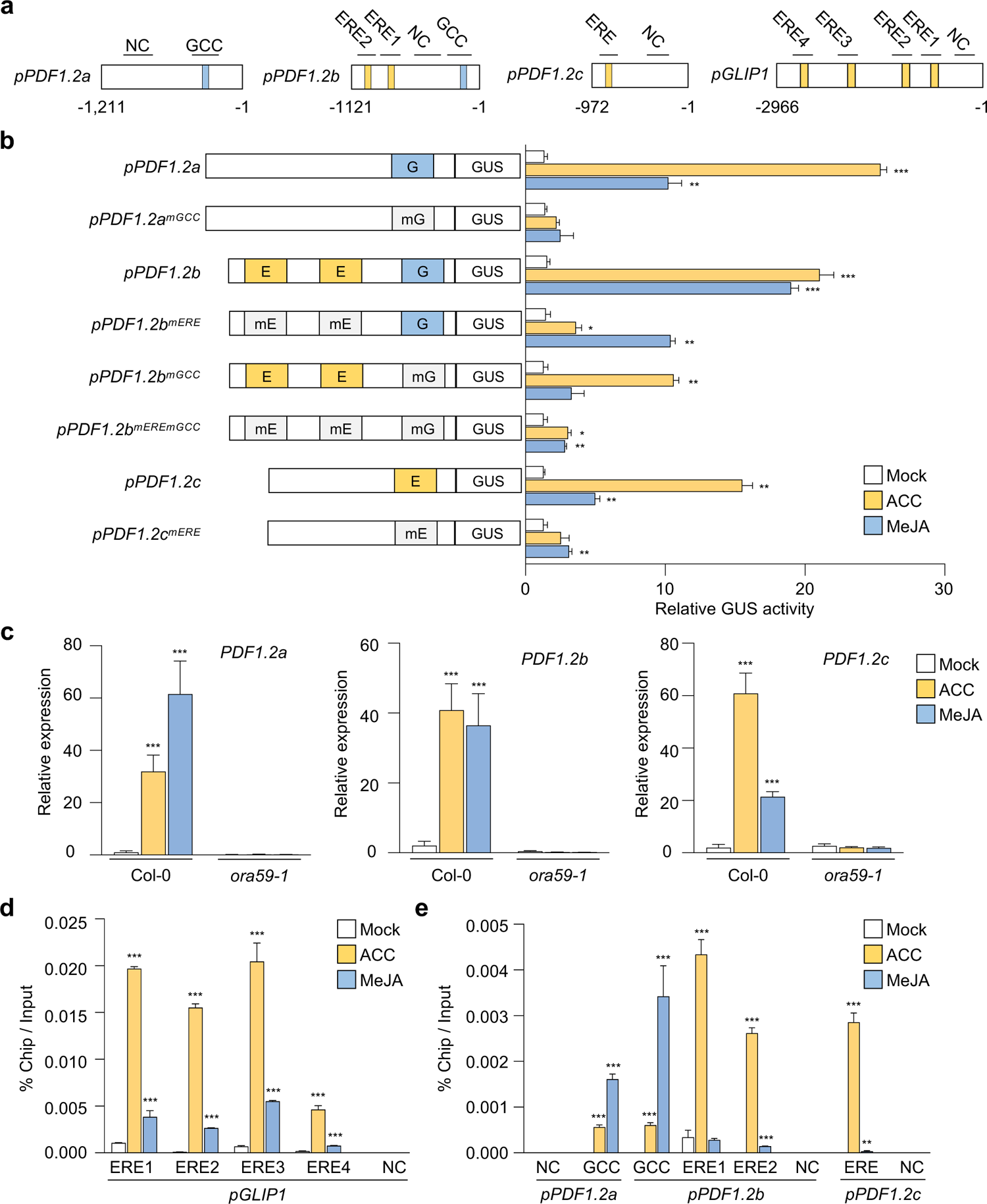
ORA59 directly binds to ERE and GCC box of *GLIP1* and *PDF1.2* promoters in ACC- and JA-dependent manners. **a**, Schematic diagram of the ERE and GCC box elements in *PDF1.2* and *GLIP1* promoters. **b**, GUS reporter assays showing the ACC- and MeJA-induced expression of the *GUS* reporter gene driven by native or ERE/GCC box-mutated *PDF1.2a*, *b*, and *c* promoters. The left panel illustrates ERE and GCC box mutations of *PDF1.2a*, *b*, and *c* promoters. E, ERE; G, GCC box; mE, mutated ERE; mG, mutated GCC box. Transfected protoplasts were treated with mock (water), ACC (200 μM), and MeJA (20 µM) for 6 h. **c**, Analysis of *PDF1.2a*, *b*, and *c* expression in ACC- and MeJA-treated plants. **d**,**e**, ChIP-qPCR analysis for *in vivo* binding of ORA59 to ERE and GCC box sequences in the *GLIP1* (**d**) and *PDF1.2* (**e**) promoters. Chromatins from ACC- and MeJA-treated *35S*:*ORA59*-*GFP* leaves were immunoprecipitated with an anti-GFP antibody. The enrichment of target element sequences is displayed as the percentage of input DNA. In **c**-**e**, six-week-old plants were treated with ACC (1 mM) and MeJA (100 µM) for 6 h. NC indicates the negative control region without ERE and GCC box sequences. Values represent means ± SD (*n* = 3 biological replicates). Asterisks indicate significant differences from mock treatment as determined by one-way ANOVA with Tukey test (\**P* < 0.05; \*\**P* < 0.01; \*\*\**P* < 0.001).

Given that ORA59 binds to ERE and GCC box in EMSA, we performed chromatin immunoprecipitation (ChIP)-qPCR analysis to examine whether ERE- and GCC box-driven transcriptional activation is induced through direct ORA59 binding to these elements in the *GLIP1* and *PDF1.2* promoters. *35S:ORA59-GFP* plants were treated with ACC and MeJA, and their extracts were used for precipitating ORA59-bound DNA fragments with an anti-GFP antibody. In the *GLIP1* promoter, all four ERE-containing fragments were enriched in ORA59 binding, and this enrichment was increased more significantly with ACC treatment than with MeJA (Fig. 6d). In addition, ORA59 binding was greatly enriched at ERE and GCC box sites of the *PDF1.2a*, *PDF1.2b*, and *PDF1.2c* promoters in response to ACC and MeJA treatments, respectively, which was consistent with the results of transcriptional activation (Fig. 6e). No binding of ORA59 was observed in negative control fragments without ERE and GCC box sequences. These results indicate that ORA59 regulates ET- and JA-responsive gene expression by binding to ERE and GCC box directly and with hormone-dependent differential preference.

### Identification of ORA59-regulated ET- and JA-responsive genes by RNA-seq analysis

Based on differential responses of ORA59 to ACC and MeJA in gene regulation, we speculated that ORA59 may regulate distinct gene sets in the ET and JA pathways. Therefore, to identify ET- and JA-responsive ORA59 downstream genes, we performed RNA-sequencing (RNA-seq) analysis using biological replicates of ACC- and MeJA-treated Col-0 and *ora59* plants (Supplementary Table 3). Differentially expressed genes (DEGs) between mock (water) and ACC/MeJA treatments were selected in Col-0 and *ora59* plants based on the cutoff (adjusted *P* (*P*adj) < 0.05, log2 fold change (|log2 FC|) ≥ 1). Col-0 had far more DEGs than *ora59* mutant, showing that 516 and 105 genes were differentially expressed in ACC-treated Col-0 and *ora59*, and 683 and 134 genes in MeJA-treated Col-0 and *ora59*, respectively (Fig. 7a). This implies that ORA59 is an essential regulator of gene expression in ET and JA responses. Considering that DEGs in *ora59* mutant are ORA59-independent, among 516 and 683 DEGs in ACC- and MeJA-treated Col-0, after subtracting 37 and 87 genes co-regulated in Col-0 and *ora59*, 479 and 596 genes were defined as ACC- and MeJA-responsive ORA59-regulated genes, respectively (Fig. 7b). It was noted that the majority of ORA59-dependent genes were upregulated in response to ACC (346 out of 479, 72.2%) and in response to MeJA (443 out of 596, 74.3%), suggesting that ORA59 primarily functions as a transcriptional activator of gene expression. The overlap between ACC- and MeJA-responsive ORA59-regulated genes was relatively small, and only 54 genes were shared (Supplementary Fig. 5).

**Figure 7.**
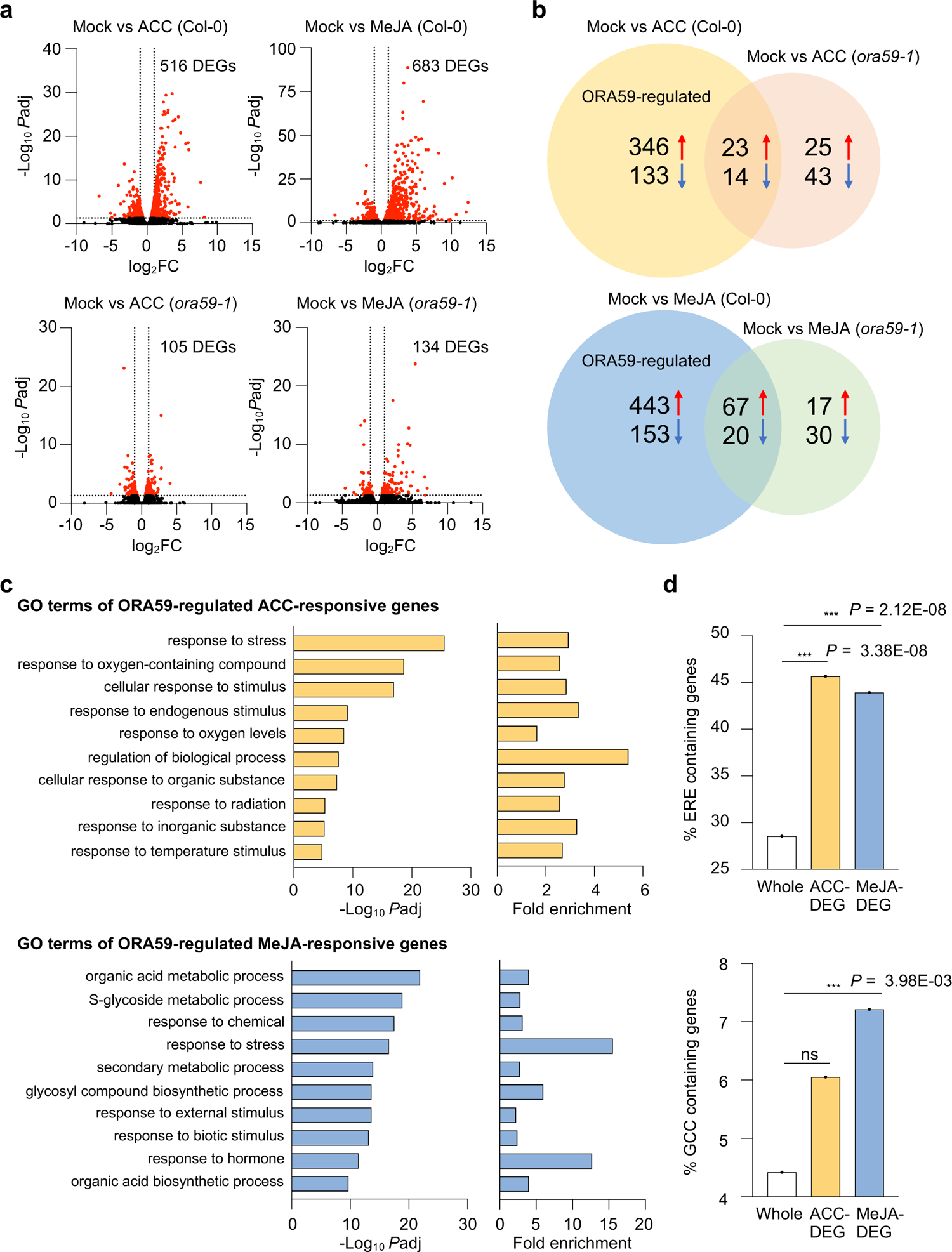
Identification ORA59-regulated ET- and JA-responsive genes by RNA-seq analysis. **a**, Volcano plots of DEGs between mock and ACC/MeJA treatments in Col-0 and *ora59* plants. Cutoff values (*P*adj = 0.05 and |log2 FC| = 1) are indicated by dashed lines. The red dots represent significantly upregulated and downregulated DEGs. **b**, Venn diagram of upregulated and downregulated DEGs between mock and ACC/MeJA treatments in Col-0 and *ora59* plants. **c**, GO enrichment analysis of ORA59-regulated DEGs. The 10 most significantly (FDR < 0.05) enriched GO terms in the Biological Process are presented for ACC- and MeJA-responsive genes. **d**, Analysis of the occurrence and enrichment of ERE and GCC box in promoters of ORA59-regulated ACC- and MeJA-responsive genes relative to whole *Arabidopsis* genes. The occurrence of ERE (AWTTCAAA) and GCC box (GCCGCC) sequences was analyzed using the regulatory sequence analysis tool (RSAT). Statistical significance of enrichment was determined by Fisher exact test (\*\*\**P* < 0.001). Whole, whole *Arabidopsis* genes; ns, not significant.

We then performed Gene Ontology (GO) enrichment analysis of ORA59-regulated DEGs, using GO Biological Process (BP) terms provided by PANTHER database (http://geneontology.org). (Supplementary Table 4). The analysis revealed that ACC-responsive DEGs are enriched in responses to stress, oxygen-containing compounds, stimulus, and oxygen levels GO BP terms, while MeJA-responsive DEGs are enriched in metabolic processes of organic acids, S-glycosides, and secondary metabolites, and in responses to stress and chemicals GO BP terms (Fig. 7c). Enriched GO BP terms indicate that ACC and MeJA regulate distinct biological processes in an ORA59 dependent manner, only co-regulating the ‘response to stress’. We then determined the occurrence and enrichment of ERE and GCC box in promoters of ORA59-regulated ACC- and MeJA-responsive genes, compared to whole *Arabidopsis* 34362 genes. Noticeably, ERE was present at a much higher frequency (28.5%) than GCC box (4.4%) in whole gene promoters (Fig. 7d). Statistically significant enrichment of ERE was observed in both ACC (Fisher exact test *P* = 3.38×10^-8^)- and MeJA (Fisher exact test *P* = 2.12×10^-8^)-responsive genes, but GCC box was only enriched in MeJA (Fisher exact test *P* = 3.98×10^-3^)-responsive genes.

### Identification of ORA59 target genes involved in disease resistance

For further functional analysis, we focused primarily on genes whose expression was increased by ACC and MeJA treatments. Among ACC- and MeJA-responsive DEGs, 63 (|log2 FC| ≥ 2) and 55 (|log2 FC| ≥ 3) upregulated genes were selected from the five most significantly enriched GO BP terms, respectively, and their expression was validated by RT-qPCR analysis (Supplementary Table 5). Towards isolating ORA59 target genes involved in the immune response, the expression of selected genes was assessed in *35S:ORA59-GFP* and *B. cinerea*–treated Col-0 plants, among which ACC-responsive eight (|log2 FC| ≥ 4) and MeJA-responsive seven (|log2 FC| ≥ 4) genes were chosen to further investigate their functions in disease resistance (Supplementary Fig. 6 and Fig. 8a).

**Figure 8.**
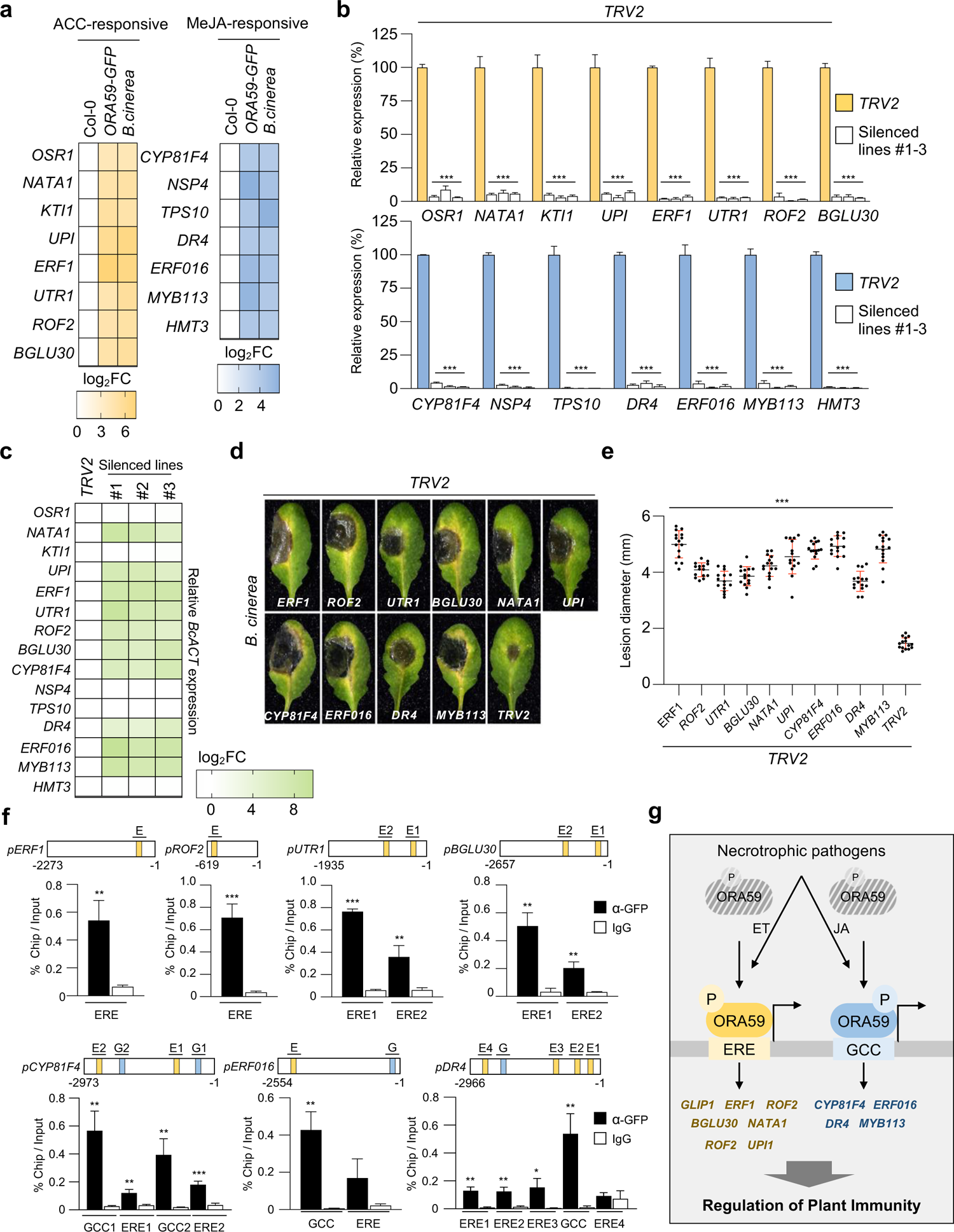
Identification of immunity-associated ORA59 target genes. **a**, Heatmap showing the transcriptional levels of the most expressed DEGs in *35S:ORA59-GFP* and *B. cinerea*-treated Col-0 plants relative to Col-0 plants. ACC-responsive eight (|log2 FC| ≥ 4) and MeJA-responsive seven (|log2 FC| ≥ 4) genes were selected for VIGS analysis. **b**, RT-qPCR analysis of the suppression of selected gene expression by VIGS. The transcript levels of selected genes in VIGS plants were determined relative to TRV2 control plants. Values represent means ± SD (*n* = 3 biological replicates). **c**, Heatmap showing the abundance of *B. cinerea actin* (*BcACT*) gene relative to *Arabidopsis tubulin 2* (*AtTU2*) in *B. cinerea*-treated TRV2 control and VIGS plants. **d**,**e**, Disease symptoms (**d**) and lesion diameters (**e**) in *B. cinerea*-treated TRV2 control and VIGS leaves. Values represent means ± SD (*n* = 15 infected leaves). **f**, ChIP-qPCR analysis for *in vivo* binding of ORA59 to ERE and GCC box sequences in ORA59 target gene promoters. Chromatins from *35S*:*ORA59*-*GFP* leaves were immunoprecipitated with an anti-GFP antibody using pre-immune IgG as a negative control. The enrichment of target element sequences is displayed as the percentage of input DNA. Values represent means ± SD (*n* = 3 biological replicates). E, ERE; G, GCC box. **g**, Model for the mechanism of ET/JA-responsive gene expression regulation by ORA59. Pathogenic infection triggers ET and JA biosynthesis, likely with different kinetic patterns, resulting in the activation of ET and JA signaling. These two hormone pathways lead to phosphorylation and stabilization of ORA59. ORA59 undergoes phosphorylation at different Ser/Thr residues in ET- and JA-dependent manners. This enhances DNA binding and transactivation activities of ORA59 with differential preference for ERE and GCC box. In this way, ET- and JA-activated ORA59 regulates different sets of genes and leads to fine-tuning of immune responses. In **c**-**e**, TRV2 control and VIGS plants were treated with 5 µl droplets of *B. cinerea* spore suspensions (5 x 10^5^ spores ml^-1^) for 2 days. Asterisks indicate significant differences from the TRV2 control (**b**,**e**) and the pre-immune IgG control (**f**) as determined by one-way ANOVA with Tukey test (\**P* < 0.05; \*\**P* < 0.01; \*\*\**P* < 0.001).

We performed tobacco rattle virus (TRV)-based virus-induced gene silencing (VIGS) in *Arabidopsis* Col-0 (Ahn et al., 2015; Burch-Smith et al., 2006). TRV2 vector (control) and VIGS constructs carrying DNA fragments of 15 genes were transformed into *Agrobacterium*, which was followed by infiltration into true leaves of seedlings. Phenotypes of VIGS plants and transcript levels of target genes were determined in rosette leaves at 19 to 21 days post-infiltration (dpi). Consistent with previous observations (Burch-Smith et al., 2006), silencing of *phytoene desaturase* (*PDS*), used as a marker of VIGS, caused photo-bleaching of leaves and reduction of *PDS* expression compared to the TRV2 control (Supplementary Fig. 7). All VIGS plants were successfully generated with more than 90% reduction in gene expression, compared to the TRV2 control, and therefore challenged with *B. cinerea* (Fig. 8b). Disease development, lesion size, and abundance of fungal actin gene were determined in the infected leaves (Fig. 8c-e). It was observed that susceptibility is increased by VIGS of 10 genes, i.e., ACC-responsive 6 and MeJA-responsive 4 genes, encoding ERF1, rotamase FKBP 2 (ROF2)/FK506-binding protein 65 (FKBP65), UDP-glucose/galactose transporter 1 (UTR1), beta glucosidase 30 (BGLU30)/dark inducible 2 (DIN2), N-acetyltransferase activity 1 (NATA1), unusual serine protease inhibitor (UPI), cytochrome P450, family 81, subfamily F, polypeptide 4 (CYP81F4), drought-repressed 4 (DR4), ERF016, and myb domain protein 113 (MYB113). Consistently, the expression of these genes was significantly reduced in *ora59* plants compared to Co-0, in response to *B. cinerea* infection, suggesting that they function downstream of ORA59 in disease resistance (Supplementary Fig. 8). Five other genes displayed no altered responses to *B. cinerea* after VIGS, perhaps because they are not related to disease resistance or it is attributed to genetic redundancy.

The isolated ORA59 target genes were scanned for the presence of ERE and GCC box in their promoters. Whereas 3 out of 10 genes had none of these elements, the other ACC- and MeJA-responsive genes contained one or more ERE and/or GCC box elements in the promoters (Fig. 8f). We carried out ChIP-qPCR analyses using *35S:ORA59-GFP* plants to determine whether ERE- and GCC box-containing genes are regulated by direct binding of ORA59 to their promoters. ChIP assays showed that ORA59 binding is enriched at ERE- and GCC box motifs of the promoters, indicating that they are direct targets of ORA59. These results suggest that ORA59 modulates immune responses to necrotrophic pathogens through regulation of direct and indirect target genes with diverse activities (Fig. 8g).

## Discussion

Phytohormone signaling and crosstalk are critical for regulating plant immune responses. In particular, JA and ET have been identified as defense signals required for resistance against necrotrophic pathogens (Dong, 1998; Pieterse et al., 2009; Zhu, 2014). Upon pathogen infection, JA and ET are synthesized rapidly and they work together, forming signaling networks which involve interactions among signaling components (Koornneef and Pieterse, 2008; Yang et al., 2015). Given that hormone signaling evokes output responses through gene regulation, JA and ET induce the expression of defense genes, such as *PDF1.2*, in a synergistic and interdependent manner (Koornneef and Pieterse, 2008; Thomma et al., 1998; Thomma et al., 1999). In this context, we observed that there is an overlap between JA- and ET-responsive genes, but on the other hand, other subsets of genes were differentially regulated by JA and ET, which was also described previously (Schenk et al., 2000).

JA/ET-mediated gene transcription typically occurs through the action of ERFs, among which ERF1 and ORA59 regulate *PDF1.2* expression by binding to the GCC box and have been considered as integrators of JA and ET signaling (Lorenzo et al., 2003; Pré et al., 2008a; Zarei et al., 2011). While ERF1 and ORA59 have been shown to regulate gene expression commonly in JA and ET pathways, questions are raised about how they respond differently to each hormone to induce JA- and ET-specific gene expression. In this study, we identified the poorly characterized ERE as an ORA59-binding *cis*-element in addition to the GCC box. EMSA, GUS reporter assays, and ChIP-qPCR analysis demonstrated that JA and ET enhance protein stability, and DNA binding and transactivation activities of ORA59 with differential preference for GCC box and ERE, respectively. While supporting this, ORA59 regulated genes of different functional categories in JA and ET pathways, as shown by RNA-seq and GO analysis. Our results provide insights into the molecular basis of how JA and ET modulate ORA59 to cooperatively and differentially regulate gene expression and to accomplish the fine-tuning of immune responses (Fig. 8g).

EIN3 functions as a positive regulator of *ERF1* and *ORA59* expression (Solano et al., 1998; Zander et al., 2012; Zhu et al., 2011). JA- and ET-mediated induction of *ERF1* and *ORA59* was abolished in *ein3 eil1* mutant, and *ORA59* promoter activity was increased by *EIN3* overexpression. In addition, EIN3 directly bound to the *ERF1* promoter, indicating that *ERF1* is the target gene of EIN3. Two different nucleotide sequences, TACAT and TTCAAA, have been identified as the EIN3-binding site (EBS) in the promoters of several EIN3-regulated genes, such as *ERF1*, *EBF2*, *protochlorophyllide oxidoreductase A* and *B* (*PORA/B*), *hookless 1* (*HLS1*), and *microRNA164* (*miR164*) (An et al., 2012; Huang et al., 2020; Konishi and Yanagisawa, 2008; Li et al., 2013; Solano et al., 1998; Zhong et al., 2009), and here they will be referred to as EBS1 and EBS2, respectively. In the course of this work, we have recognized that the EBS2, TTCAAA, is part of the ERE sequence AWTTCAAA, and this is especially true for the EBS2 in *PORB* and *HLS1* promoters. Therefore, we wonder whether EIN3-regulated genes containing the EBS2 in their promoters, which overlaps with the ERE, are also targets of ORA59, and whether ORA59 is implicated in other cellular processes, such as light signaling and seedling development. It may be possible that EIN3 and ORA59 share and co-regulate certain target genes, which is supported by the evidence that EIN3 and ORA59 proteins interact together (He et al., 2017).

In our RNA-seq analysis, ERF1 was isolated as an ORA59-regulated ET-responsive gene. In addition to the previously identified EBS2 (Solano et al., 1998), the *ERF1* promoter has a separate ERE, to which ORA59 directly bound as determined by ChIP analysis, implying that ORA59 is the upstream regulator of *ERF1*. Conversely, a previous study showed that *ORA59* expression is largely increased in *ERF1*-overexpressing plants (Van der Does et al., 2013). Given that the *ORA59* promoter contains both ERE and GCC box, ORA59 and ERF1 may activate each other via a positive feedback loop. Furthermore, ERF1 bound to another stress–responsive element DRE/CRT during abiotic stress responses, as shown in a previous study (Cheng et al., 2013). This and our data suggest that ERFs, including ERF1 and ORA59, may bind to distinct types of *cis*-elements, depending on hormone and stress stimuli. On the other hand, studies have shown that other transcription factors, TGA2/4/6 (class II TGAs) and WRKY33, positively regulate *ORA59* expression in response to ACC and *B. cinerea* infection through binding to the TGA binding site TGACGT and the W-box TTGAC(C/T) in the *ORA59* promoter, respectively (Birkenbihl et al., 2012; Zander et al., 2014). Further investigation is needed on how ET/JA-regulated EIN3, ORA59, and ERF1, and other types of transcription factors, such as TGAs and WRKYs, interact and coordinately regulate gene expression in the transcriptional and protein interaction networks.

Protein phosphorylation regulates the function of transcription factors by modulating DNA binding, transcriptional activity, protein stability, cellular localization, and protein-protein interactions. Many reports provide evidence that ERFs are regulated by phosphorylation (Huang et al., 2016; Licausi et al., 2013; Phukan et al., 2017). Phosphorylation of the tomato ERF Pti4 by Pto kinase enhanced Pti4 binding to the GCC box, increasing the expression of GCC box-containing *PR* genes (Gu et al., 2000). Mitogen-activated protein kinase (MAPK/MPK) cascades have been implicated in ERF phosphorylation. When phosphorylated by blast and wound-induced MAPK1 (BWMK1), the rice ET-responsive element-binding protein 1 (OsEREBP1) showed enhanced DNA binding activity to the GCC box, and concomitantly, the increased GCC box-driven transcription (Cheong et al., 2003). Transactivation by the tobacco NtERF221 (originally designated as ORC1) was positively affected by a MAPK kinase, JA-factor stimulating MAPKK1 (JAM1) (De Boer et al., 2011). The *Arabidopsis* ERF6 served as an MPK substrate, and its protein stability and nuclear localization were increased by MPK3/MPK6-mediated ERF6 phosphorylation (Meng et al., 2013; Wang et al., 2013). In this study, we showed that ORA59 phosphorylation is elevated in plants treated with either ACC or MeJA. ORA59 activated through ACC and JA signals had differential preferences for ERE and GCC box, in addition to enhanced DNA binding, which was eliminated by phosphatase-mediated dephosphorylation of ORA59. Likewise, ORA59 proteins which accumulated in MG132-treated plants showed a similar level of binding to these elements. These results suggest that phosphorylation regulates both affinity and specificity of ORA59 for DNA sequences. Furthermore, recombinant ORA59 proteins with deletion of the N- and C-terminal parts showed differential GCC box- and ERE-binding activities, suggesting that ORA59 may form distinct structures with different affinities for ERE and GCC box, and this may be regulated by hormone-dependent phosphorylation events (Fig. 8g). Therefore, it is important to investigate whether ORA59 phosphorylation occurs in ET- and JA-dependent ways, and whether it modulates the structure and activity of ORA59. A combination of ACC and MeJA treatments further increased the level of phosphorylation, but not that of DNA binding activity of ORA59, suggesting that ORA59 phosphorylation may be involved in ET and JA synergy at the level of transcriptional activation, e.g., interaction with other cofactors/transcription factors and transcription machinery components.

Gene expression, VIGS, and ChIP analysis led to the identification of direct target genes of ORA59, *ERF1*, *ROF2*/*FKBP65*, *UTR1*, *BGLU30*/*DIN2* as ACC-responsive genes, and *CYP81F4*, *DR4*, and *ERF016* as MeJA-responsive genes, and indirect target genes, *NATA1*, *UPI*, and *MYB113*. These ORA59 target genes are clustered into four functional groups. First, *ERF1*, *ERF016*, and *MYB113* encode transcription factors, which are involved in the regulation of defense gene expression. ERF1 has been well characterized to enhance *PDF1.2* expression and disease resistance (Zarei et al., 2011). ERF016 bound to the GCC box of the *PDF1.2* promoter and *erf016* mutants displayed a significant increase in susceptibility to *B. cinerea* (Hickman et al., 2017; Ou et al., 2011). *MYB113* expression was much reduced in *ora59* mutants, suggesting that MYB113 functions downstream of ORA59 (Zander et al., 2014). Second, *DR4* and *UPI* encode protease inhibitors implicated in resistance to necrotrophic fungi (Brodersen et al., 2006; Gosti et al., 1995; Laluk and Mengiste, 2011). Third, *BGLU30*/*DIN2*, *CYP81F4*, and *NATA1* function in secondary metabolic pathways. *BGLU30*/*DIN2*, encoding a β-glucosidase, and *CYP81F4*, encoding a cytochrome P450 monooxygenase, showed activities associated with glucosinolate metabolism (Morikawa-Ichinose et al., 2020; Pfalz et al., 2011; Zhang et al., 2020). Glucosinolates and their breakdown products function in defense against pathogens (Bednarek, 2012), supporting the possibility that BGLU30 and CYP81F4 may play a role in plant immunity. NATA1 was identified as an acetyltransferase that acetylates ornithine and putrescine in response to coronatine, JA, and *P. syringae* infection (Adio et al., 2011; Lou et al., 2016). Fourth, *ROF2*/*FKBP65*, encoding a peptidyl-prolyl cis-trans isomerase, and *UTR1*, encoding a nucleotide sugar transporter, are involved in protein folding and endoplasmic reticulum (ER) quality control processes. Knockout or overexpression of *ROF2*/*FKBP65* decreased or increased resistance against *P. syringae*, respectively (Pogorelko et al., 2014). UTR1, required for the transport of UDP-glucose into the ER, may be involved in plant immunity, because proper folding of immune receptors and PRRs relies on the ER quality control system (Eichmann and Schäfer, 2012; Reyes et al., 2006). Further studies on the functions of ORA59 target genes will improve our understanding of the ET-JA signaling network and involving components in the regulation of plant immunity.

## Methods

### Plant materials and growth conditions

*Arabidopsis thaliana* (ecotype Columbia, Col-0) plants were grown at 23°C under long-day conditions in a 16-h light/8-h dark cycle. The mutant lines used in this study are *glip1-1* (Oh et al., 2005), *ein2-1* (Roman and Ecker, 1995), *ein3-1eil1-1* (Alonso et al., 2003), *ora59* (CS_405772), and *coi1* (SALK_095916). Homozygous lines were selected by PCR and sequence analysis using gene-specific primers (Supplementary Table 6). To generate *35S:ORA59-GFP* plants, the *ORA59* coding region was cloned into the pCHF3-GFP binary vector under the control of the CaMV 35S promoter. To generate *pGLIP1:GUS* and *pGLIP1^mEREs^:GUS* plants, the *GLIP1* promoter region (−1 to −2966 bp) was amplified from *Arabidopsis* gDNA by PCR and cloned into the pBI121 vector containing a *GUS* gene. ERE mutations in the *GLIP1* promoter were generated by site-directed mutagenesis using primers in Supplementary Table 6. For *pGLIP1:GLIP1-GFP* and *pGLIP1^mEREs^:GLIP1-GFP* plants, the *GLIP1* coding region was cloned into the pCAMBIA1300 vector containing a *GFP* gene, and then *pGLIP1* and *pGLIP1^mEREs^* were inserted upstream of *GLIP1-GFP* in the pCAMBIA1300 vector. The constructs were transformed into *Agrobacterium tumefaciens* GV3101 and then introduced into Col-0 and *glip1-1* plants using the floral dip method (Clough and Bent, 1998).

### Plant treatments

For pathogen infection, *B. cinerea* and *A. brassicicola* were grown on potato dextrose agar plates for 2 weeks, and their spores were harvested and incubated in half-strength potato dextrose broth for 2 h prior to inoculation as previously described (Broekaert et al., 1990). Six-week-old leaves were inoculated with 5 µl droplets of spore suspensions (5 x 10^5^ spores ml^-1^). Fungal growth was assessed by qPCR for the abundance of *A. brassicicola cutinase A* (*AbCUTA*) and *B. cinerea actin* (*BcACT*) genes relative to *Arabidopsis tubulin 2* (*AtTU2*). Lesion size was determined by measuring the diameter of the necrotic area. For chemical treatments, 6-week-old plants were sprayed with 0.01% Silwet L-77 containing 1 mM SA, 1 mM ACC, 100 µM MeJA, 50 µM MG132, and 100 µM CHX or incubated with 10 ppm ET in hydrocarbon-free air. The treated plants were maintained at 100% humidity for the indicated times.

### Transient expression assays

For transient assays in *Arabidopsis* protoplasts, effector and reporter constructs were generated. For effector constructs, coding regions of *ORA59*, *RAP2.2*, and *CPL3* were amplified from the *Arabidopsis* cDNA library by PCR and cloned into the pUC18 vector for the expression of hemagglutinin (HA)-tagged proteins in protoplasts (Cho and Yoo, 2011). For gene promoter-reporter constructs, promoter regions of *GLIP1*, *PDF1.2a*, *PDF1.2b*, and *PDF1.2c* were amplified from *Arabidopsis* gDNA by PCR and cloned into the pBI221 vector containing a *GUS* gene. Mutations of ERE and GCC box elements in the promoter regions were generated by site-directed mutagenesis using primers in Supplementary Table 6. For synthetic promoter-reporter constructs, four copies of the native ERE or GCC box and four copies of respective mutated versions were fused with the minimal *GLIP1* promoter (−1 to −122 bp) and cloned into the pBI221 vector containing a *GUS* gene. *Arabidopsis* mesophyll protoplasts were isolated and transfected as previously described (Yoo et al., 2007). Isolated protoplasts (2 x 10^4^) were transfected with a reporter DNA (20 μg) alone or together with an effector DNA (20 μg). GUS activity was measured fluorometrically using 4-methylumbelliferyl-β-D-glucuronide as substrate. The firefly luciferase (LUC) expressed under the control of the CaMV 35S promoter was used as an internal control, and the activity was measured using the luciferase assay system (Promega). Relative GUS activities were normalized with respect to the LUC activity.

### Histochemical GUS staining

GUS staining was performed as previously described (Lee et al., 2017a). Rosette leaves were incubated in a staining buffer (50 mM sodium phosphate, pH 7.0, 0.5 mM K_3_Fe(CN)_6_, 0.5 mM K_4_Fe(CN)_6_, 10 mM EDTA, and 0.2% Triton X-100) containing 4 mM 5-bromo-4-chloro-3-indolyl-b-D-glucuronide (X-Gluc) for 16 h at 37°C. Stained leaves were cleared by several washes with 70% ethanol.

### Yeast one-hybrid (Y1H) screening

Y1H screening was performed as previously described (Welchen et al., 2009). To obtain a yeast strain carrying the ERE sequence in front of the *HIS3* reporter gene, three tandem repeats of the ERE were cloned into the pHIS3-NX vector, and the 3xERE-*HIS3* cassette was cloned into the pINT vector, which confers resistance to the antibiotic G418. The clone in pINT1 was introduced into the yeast strain Y187. Alternatively, the 3xERE was placed in front of the *lacZ* reporter gene contained in the pLacZi vector (Clontech). Transcription factors interacting with the ERE sequence were identified using a DNA library carrying a 1050 *Arabidopsis* transcription factor ORFeome collection in the prey vector pDEST22 (Invitrogen). For Y1H screening, plasmid DNA from the library (10 μg) was introduced into yeast and a total of 2 x 10^6^ transformants were plated on SD-Trp-His medium containing 0.2 mM 3-AT. The resulting putative positive clones were streaked on fresh SD-Trp-His + 0.2 mM 3-AT medium to purify colonies. The plasmid DNAs containing ORFs were rescued and retransformed into yeast for confirmation.

### Protein expression and purification

The full-length coding regions of *ORA59*, *RAP2.2*, *CPL3*, and the truncated regions of *ORA59* were PCR-amplified using gene-specific primers (Supplementary Table 6). The PCR products were cloned into the pMAL-x2X vector to generate proteins fused to the N-terminal maltose-binding protein (MBP) and His-tag. *Escherichia coli* BL21(DE3) pLysS cells were transformed with the constructs and cultured at 28°C. Protein expression was induced by the addition of 0.3 mM IPTG for 3 h at 28°C. The MBP/His-tagged proteins were purified using Ni^2+^-NTA agarose (Qiagen) according to the manufacturer’s instructions.

### Nuclear extraction

Five-week-old leaves were ground in liquid nitrogen and incubated in a nuclear extraction buffer (20 mM PIPES-KOH, pH 7.0, 10 mM KCl, 1.5 mM MgCl_2_, 0.3% Triton X-100, 5 mM EDTA, 1 mM DTT, 1 M 2-methyl-2,4-pentandiol, 1 mM NaF, 1 mM Na_3_VO_4_, and protease inhibitor cocktail) on ice for 10 min. The material was filtered through one layer of Miracloth and spun at 1,000 g for 10 min at 4°C. After removing the supernatant, the pellet was resuspended in a buffer (20 mM PIPES-KOH, pH 7.0, 10 mM MgCl_2_, 1% Triton X-100, 1 mM DTT, 0.5 M hexylene glycol (2-methyl-2,4-pentandiol), 1 mM NaF, and 1 mM Na_3_VO_4_), incubated on ice for 10 min, and then centrifuged at 1,000 g for 10 min at 4°C. To extract nuclear proteins, isolated nuclei were resuspended in an extraction buffer (20 mM HEPES, pH 8.0, 300 mM NaCl, 1 mM MgCl_2_, 0.2 mM EDTA, 10% glycerol, 1% Triton X-100, 0.1% NP-40, 1 mM DTT, 1 mM NaF, 1 mM Na_3_VO_4_, and protease inhibitor cocktail), incubated with rotation for 30 min at 4°C, and then centrifuged at 15,000 g for 20 min at 4°C. The supernatant was collected, and the protein concentration was determined before use. For phosphatase treatment, phosphatase inhibitors (NaF and Na_3_VO_4_) were excluded from the extraction buffer, and extracted nuclear proteins were treated with lambda protein phosphatase (NEB) according to the manufacturer’s instruction.

### Electrophoretic mobility shift assay (EMSA)

EMSA was performed using the LightShift Chemiluminescent EMSA kit (Thermo Scientific). Biotin-labeled oligonucleotides were synthesized by Macrogen (Korea). Purified proteins or nuclear extracts were incubated in 20 fM biotin-labeled oligonucleotide probes in 15 μl of a binding buffer (10 mM Tris-HCl, pH 7.5, 40 mM KCl, 3 mM MgCl_2_, 1 mM EDTA, 10% glycerol, 1 mM DTT, and 3 μg poly(dI-dC)) for 30 min at room temperature (purified proteins) or at 4°C (nuclear extracts). The samples were resolved on 5% polyacrylamide (75:1 acrylamide:bis-acrylamide) gels. In the competition assay, purified ORA59 proteins were incubated with the indicated excess amounts of oligonucleotide competitors for 15 min before the addition of biotin-labeled probes.

### Immunoblotting and immunoprecipitation

For immunoblotting, proteins were separated on 10-12% SDS polyacrylamide gels by SDS-gel electrophoresis and electro transferred onto nitrocellulose membranes. Membranes were incubated with anti-GFP (sc-9996, Santa Cruz Biotechnology), anti-Actin (ab197345, Abcam), anti-Histone H3 (ab1791, Abcam), and anti phospho Ser/Thr (ab17464, Abcam) antibodies. For immunoprecipitation, nuclear pellets were lysed in hypotonic buffer (20 mM HEPES, pH 7.9, 20 mM KCl, 1.5 mM MgCl_2_, and 25% glycerol) and high-salt buffer (20 mM HEPES, pH 7.9, 800 mM KCl, 1.5 mM MgCl_2_, 25% glycerol, and 1% NP-40) supplemented with protease inhibitor cocktail and incubated with rotation at 4°C. Lysates were cleared by centrifugation and incubated with an anti-GFP antibody for 2 h at 4 °C. After an additional 2 h incubation with Protein G Agarose (20399, Thermo Scientific), beads were washed with wash buffer (20 mM Tris-HCl, pH 7.9, 150 mM KCl, 20% glycerol, 0.1mM EDTA, and 0.1% NP-40) and bound proteins were eluted with 2x sample buffer (100 mM Tris-HCl, pH 6.5, 20% glycerol, 4% SDS, 200 mM DTT, and 3 mM bromophenol blue). Immunoblot bands were visualized using the enhanced chemiluminescence system (Amersham Biosciences).

### Gene expression analysis

Total RNAs were extracted using TRIzol reagent and reverse-transcribed into cDNAs using the PrimeScript RT reagent kit (TaKaRa). RT-qPCR was performed using KAPA SYBR FAST qPCR master mix (Kapa Biosystems) with gene-specific primers (Supplementary Table 6) on a LightCycler 480 system (Roche) according to the manufacturer’s protocol. For transcript normalization, *Actin1* was used as a reference gene. Data were analyzed using LC480Conversion and LinRegPCR software (Heart Failure Research Center).

### RNA sequencing data analysis

Total RNAs were extracted from leaves using RNeasy Plant Mini kit (Qiagen). The amount of RNAs was measured using Nanodrop (Thermo Scientific) and the quality was assessed using Bioanalyzer (Agilent Technologies) with an RNA Integrity Number (RIN) value ≥ 8. The RNA-seq libraries were prepared using the TruSeq RNA preparation kit V2 kit following the manufacturer’s instructions. The 150-bp paired-end sequencing reads were generated on the Illumina NextSeq 550 System instrument platform. The low-quality base (base quality score < 20) in the last position of the reads was trimmed and high-quality sequencing reads were subsequently aligned onto the *A. thaliana* reference genome (TAIR10) using HISAT2 (Kim et al., 2019). The raw number of reads mapped onto each transcript was quantified using StringTie (Pertea et al., 2015) and the counts per transcript were normalized based on the library size by DESeq2 (Love et al., 2014). The batch effect among samples was estimated by PCA and corrected by limma (Ritchie et al., 2015). Statistically significant DEGs were tested based on a negative binomial distribution using a generalized linear model. Enriched GO terms for DEGs were determined using the statistical overrepresentation test in PANTHER (http://geneontology.org). Gene lists were compared to all *Arabidopsis* genes in PANTHER using the GO BP dataset and binomial test with FDR correction.

### Chromatin immunoprecipitation (ChIP)

ChIP assays were performed as described previously (Lee et al., 2017b). Five-week-old *35S:ORA59-GFP* leaf tissues were fixed with 1% formaldehyde under vacuum, washed, dried, and ground to a fine powder in liquid nitrogen. The powder was suspended in M1 buffer (10 mM sodium phosphate, pH 7.0, 0.1 M NaCl, 1 M 2-methyl 2,4 pentanediol, 10 mM β-mercaptoethanol, and protease inhibitor cocktail). Nuclei were isolated from the filtrate by centrifugation at 1000 g for 20 min at 4°C and washed with M2 buffer (10 mM sodium phosphate, pH 7.0, 0.1 M NaCl, 1 M 2-methyl 2,4 pentanediol, 10 mM β-mercaptoethanol, 10 mM MgCl_2_, 0.5% Triton X-100, and protease inhibitor cocktail) and M3 buffer (10 mM sodium phosphate, pH 7.0, 0.1 M NaCl, 10 mM β-mercaptoethanol, and protease inhibitor cocktail). The crude nuclear pellet was resuspended in sonication buffer (10 mM sodium phosphate, pH 7.0, 0.1M NaCl, 0.5% Sarkosyl, and 10 mM EDTA) and sonicated to obtain DNA fragments. The fragmented chromatin was transferred to IP buffer (50 mM HEPES, pH 7.5, 150 mM NaCl, 5 mM MgCl_2_, 10 μM ZnSO4, 1% Triton-X 100, and 0.05% SDS). The pre-cleared chromatin was incubated with IgG or GFP antibody (A11122, Thermo Scientific) for 2 h at 4°C. After an additional overnight incubation with Protein A Sepharose (20333, Thermo Scientific), beads were incubated in elution buffer (0.1M glycine, pH 2.8, 0.5 M NaCl, 0.05% Triton X-100). Primers for qPCR are listed in Supplementary Table 6.

### Virus-induced gene silencing (VIGS)

VIGS was performed as previously described (Burch-Smith et al., 2006). Coding regions of target genes were amplified and cloned into the pTRV2 vector. The constructs were transformed into *Agrobacterium tumefaciens* strain GV3101, which was cultured in LB media containing 10 mM MES-KOH (pH 5.7), 200 μM acetosyringone, 50 mg l^-1^ gentamycin, and 50 mg l^-1^ kanamycin overnight at 28°C. *A. tumefaciens* cells were harvested, adjusted to an OD600 of 1.5 in infiltration media (10 mM MES-KOH, pH 5.7, 10 mM MgCl_2_, and 200 μM acetosyringone), and infiltrated into leaves of *Arabidopsis* seedlings at 15-17 days after germination. After 19-21 days, VIGS plants were treated with pathogens and silencing effects were verified in the systemic leaves by gene-specific primers (Supplementary Table 6).

### Statistical analysis

Statistical analyses were performed using GraphPad Prism (v. 8.0). Significant differences between experimental groups were analyzed by one-way ANOVA with Tukey’s HSD test or unpaired Student’s *t* test for multiple comparisons or single comparisons, respectively. Detailed information about statistical analysis is described in the figure legends. Statistical significance was set at *P* < 0.05. All experiments were repeated 3-5 times with similar results.

## Acknowledgments

We are grateful to H. S. Pai (Yonsei University, Korea) for technical help with VIGS. This work was supported by a Korea University grant, a Next-Generation BioGreen 21 Program (SSAC, PJ013202) from the Rural Development Administration and a National Research Foundation of Korea (NRF) grant (2018R1A5A1023599, SRC) from the Korean government (MSIP). J.C.H. acknowledges support from a NRF grant (2020R1A6A1A03044344) from the Korean government (MSIP).

## Author contributions

O.K.P. conceived and directed the project. Y.N.Y. and O.K.P. designed the research. Y.N.Y. performed most of the experiments. Y.K. performed protein purification and EMSA analysis. H.K. conducted ChIP-qPCR analysis. D.S.L. and M.H.L. participated in EMSA and ChIP-qPCR analysis. S.J.K. generated transgenic and VIGS lines. K.M.C., Y.K., S.J.K., and J.C. conducted RNA-seq analysis. S.Y.K. and J.C.H. participated in Y1H screening experiments. Y.N.Y. and O.K.P. analyzed the data and wrote the manuscript. All authors contributed to the review and editing of the manuscript.

